# Resting behaviour of malaria vectors in a highland and a lowland site of western Kenya: Implication on malaria vector control measures

**DOI:** 10.1101/815175

**Authors:** Maxwell G. Machani, Eric Ochomo, Fred Amimo, Jackline Kosgei, Stephen Munga, Guofa Zhou, Andrew K. Githeko, Guiyun Yan, Yaw A. Afrane

## Abstract

**Background:** Understanding the interactions between increased insecticide resistance in field malaria vector populations and the subsequent resting behaviour patterns is important for planning adequate vector control measures in a specific context and sustaining the current vector interventions. The aim of this study was to investigate the resting behavior, host preference and infection with *Plasmodium falciparum* sporozoites by malaria vectors in different ecological settings of western Kenya with different levels of insecticide resistance.

**Methods:** Indoor and outdoor resting *Anopheline* mosquitoes were sampled during the dry and rainy seasons in Kisian (lowland site) and Bungoma (highland site), both in western Kenya. WHO tube bioassay was used to determine levels of phenotypic resistance of first generation offspring (F1 progeny) of malaria vectors resting indoors and outdoors to deltamethrin. PCR-based molecular diagnostics were used for mosquito speciation, genotype for resistance mutations and to determine specific host blood meal origins. Enzyme-linked Immunosorbent Assay (ELISA) was used to determine mosquito sporozoite infections.

**Results:** Overall, 3,566 female *Anopheles* mosquitoes were collected with *Anopheles gambiae* s.l [In Bungoma, *An. gambiae s.s* (90.9%), *An arabiensis* (7.6%) and in Kisian, *An. gambiae s.s* (38.9%), *An. arabiensis* (60.2%)] being the most abundant species (74.7%) followed by *An. funestus* s.l (25.3%). The majority of *An. gambiae* s.l (85.4 and 58%) and *An. funestus* (96.6 and 91.1%) were caught resting indoors in Bungoma and Kisian respectively.Vgsc-1014S was observed at a slightly higher frequency in *An. gambiae s.s* hereafter(*An. gambiae*) resting indoor than outdoor (89.7 vs 84.6% and 71.5 vs 61.1%) in Bungoma and Kisian respectively. For *An. arabiensis*, Vgsc-1014S was 18.2% indoor and outdoor (17.9%) in Kisian. In Bungoma, the Vgsc-1014S was only detected in *An. arabiensis* resting indoors with a frequency of 10%. The Vgsc-1014F mutation was only present in *An. gambiae* resting indoors from both sites, but at very low frequencies in Kisian compared to Bungoma (0.8 and 9.2% respectively. In Bungoma, the sporozoite rates for *An. funestus*, *An. gambiae*, and *An. arabiensis* resting indoors were 10.9, 7.6 and 3.4 % respectively. For outdoor resting, *An. gambiae* and *An. arabiensis* in Bungoma, the sporozoite rates were 4.7 and 2.9 % respectively.Overall, in Bungoma, the sporozoite rate for indoor resting mosquitoes was 8.6% and 4.2% for outdoors. In Kisian the sporozoite rate was 0.9% for indoor resting *An. gambiae.* None of the outdoor collected mosquitoes in Kisian tested positive for sporozoite infections.

**Conclusion:** The study reports high densities of insecticide-resistant *An. gambiae* and *An. funestus* resting indoors and the persistence of malaria transmission indoors with high entomological inoculation rates (EIR) regardless of the use of Long-lasting insecticidal nets (LLINs). These findings underline the difficulties of controlling malaria vectors resting and biting indoors using the current interventions. Supplemental vector control tools and implementation of sustainable insecticide resistance management strategies are needed in western Kenya.

## Background

Malaria still remains a major public health concern in sub-Saharan Africa, responsible for an estimated 219 million cases and 435,000 deaths despite the massive investments in scaling-up indoor anti-vector interventions [1]. Remarkable advances in the fight against malaria have been achieved within the past decade mainly through the massive scale-up of long-lasting insecticide-treated nets (LLINs) and indoor residual spraying (IRS) in many localities [2, 3]. Despite the increased efforts, it is worrying that no significant progress has been made in reducing global malaria cases in the year 2015–2017 period [1], with some regions in sub-Saharan Africa, previously reported to experience a resurgence of malaria including western Kenya [4]. This transmission recurrence is partly attributed to the emergence of insecticide resistance and behavioural modification that have arisen as an adaptation by mosquitoes in response to high use of insecticides for vector control [5, 6]. All these factors have the potential to weaken malaria control programs thus posing a serious threat in the fight against malaria [7].

The current vector control interventions take advantage of susceptible mosquito behaviors. These interventions are based on the observation that malaria vectors prefer to bite humans indoors late at night and often rest inside houses after blood feeding hence, they will be exposed to sufficient levels of insecticides which will either kill them or reduce their longevity thus affecting their vectorial capacity [1]. In sub-Saharan Africa insecticide-treated net (ITN) ownership is estimated to have increased from 3% in 2000 to 83% in the period of 2015-2017[1]. In Kenya, the government rolled out the universal bed net programme where every two persons in a household were provided with a free ITN. The ITN ownership rose from 12.8% in 2004 to over 80% in 2015[8, 9] The increased use of indoor interventions may pose stress on the indoor feeding and resting of malaria vectors leading to either behavioural defense [10] or physiological defense [5].

Malaria vectors have been shown to adapt to changing environment due to either behavioural avoidance or selection of mutations and recombination that favour their survival in the presence of insecticides threatening the efficacy of the current indoor-based vector control tools [5, 11] and the resulting increase in residual transmission [12]. Insecticide resistance is common in sub-Saharan Africa with some regions reporting resistance to all classes of insecticides [13, 14]. In Kenya, the target site and metabolic resistance mechanisms play a major role in pyrethroid resistance [15, 16]. The primary malaria vectors in Kenya belong to *An. gambiae* complex and *An. funestus* group due to their anthropophilic and endophilic behaviours that makes them be more efficient in malaria transmission [17–19]. With the scale-up of indoor-based vector control tools mosquitoes have changed behaviours; some are biting and resting indoors whilst others have changed to prefer biting bite outdoors. Behavioral modifications including changes in biting time and location [20–22], changes in host choice and shift from endophilic (i.e. resting in houses) to exophilic (i.e. resting outdoors) behavior have been associated with long-term use of insecticide-based interventions [23, 24]. Insecticide resistance could make mosquitoes respond numerically and behaviorally to alter vector populations as well as resting and feeding behaviors. Knowledge of the resting habits of resistant vectors and their feeding preference may predict the intensity of malaria transmission. It is hypothesized that insecticide-resistant malaria vectors could bite and rest indoors in the presence of interventions whilst susceptible ones bite and rest outdoors. Additionally, behaviors of malaria vectors have been shown to differ on small geographical scales, further complicating malaria elimination efforts[25]. Understanding how the resting habits of malaria vectors change in response to current indoor-based vector control interventions is important for sustaining vector control. These behavioral modifications and physiological resistance in most of the malaria vectors have been shown to contribute to maintaining malaria transmission [11, 26].

In order to improve vector control intervention strategies, it is crucial to characterize the behavioural patterns of each species of a particular vectorial system in their specific settings over time and in a range of environmental changes, especially with increasing pyrethroid resistance. The objective of this study was to investigate the species diversity of malaria vectors, their resting behavior, and the distribution of infections in two ecological settings of western Kenya with different levels of insecticide resistance. This information could provide a better understanding of the interactions between increased insecticide resistance phenotypes in field malaria vector population and the subsequent resting behaviour patterns in the presence of the current indoor intervention.

## Methods

### Study site

The study was carried out in Kisian (00.02464°S, 033.60187°E, altitude 1,280–1,330 m above sea level), Kisumu county and Bungoma (00.54057°N, 034.56410°E, altitude 1386-1,545 m above sea level) in Bungoma County, all in western Kenya. Kisian is located in the lowland area around Lake Victoria in western Kenya. A*n. gambiae sensu stricto (*s.s.), *An. arabiensis* and *An. funestus* are the main vectors of malaria in this region [16, 27]. Bungoma County is located in the malaria epidemic-prone highland area with the long (May - July) and short (October-November) rainy seasons and year-to-year variation. Both sites are endemic for *Plasmodium falciparum* malaria. The malaria vector population in both sites include *An. gambiae* and *An. arabiensis* and *Anopheles funestus* [15, 16]

### Mosquito sampling

Mosquitoes were sampled during the long dry season (February-March) and the long rainy season (May-July) in 2018. Indoor resting malaria vectors were sampled using pyrethrum spray catches (PSCs) and mouth aspirators in sixty (60) randomly selected houses from 06:00 to 09:00 h [22]. Outdoor resting mosquitoes were collected from pit shelters constructed according to Muirhead-Thomson’s method [28], clay-pots and by the use of Prokopack aspirators (John W Hock, Gainesville, FL, USA) from outdoor kitchens and evening outdoor human resting points. *Anopheline* mosquitoes were sorted morphologically according to the identification keys described by [29]. Female mosquitoes were further classified according to their gonotrophic status. Mosquitoes from each collection method were stored in separately labeled vials and preserved by desiccation.

Some of the collected indoor and outdoor resting mosquitoes that were either blood fed or gravid from the two sites were kept in paper cups covered with moistened cotton towels and transported to the insectary at Kenya medical research institute in Kisumu. Gravid *An. gambiae* s.l and *An. funestus* s.l females were provided with oviposition cups. Eggs laid were allowed to hatch in spring water in small trays and larvae reared on a mixture of tetramin (fish food) and brewer’s yeast provided daily under controlled standard insectary conditions with a temperature range of 26± 2°C and 70% to 80% relative humidity. Emerging adults were provided with a 10% sugar solution until ready to be used for bioassay tests.

### WHO resistance bioassays

To assess susceptibility or resistance of F1 progeny of mosquitoes caught from different locations(indoor and outdoor) and study sites, emerging female adults aged 2-5 days were exposed to 0.05% deltamethrin following the standard WHO tube test protocol[30] for 1 h. The knockdown time (KDT) of females was reported every 10 min during the 60 min exposure period. After the 1 h exposure, surviving mosquitoes were transferred to recovery tubes and provided with 10% sucrose for 24 h holding period. Mosquitoes alive 24 h after the 60-min insecticide-exposure time were classified as resistant. Mortality was scored after the 24 h recovery period.

### *Anopheline* Species discrimination

Sibling species of the *An. gambiae* and *An. funestus* complexes were distinguished using conventional PCR [31, 32]. DNA was extracted from mosquito legs and wings using ethanol precipitation method [33].

### Genotyping for kdr mutations

DNA was extracted from adult *An. gambiae* and *An. arabiensis* mosquitoes as earlier described [31]. Real-time (RT) PCR was used to detect mutations at amino acid position 1014 of the voltage-gated sodium channel (Vgsc) using a modification of the protocol by Bass et al. [34]. Samples were genotyped for both *Vgsc*-1014S and 1014F kdr alleles.

### Detection of blood meal sources using polymerase chain reaction (PCR)

The abdominal section of blood-fed *Anopheles* mosquitoes were cut transversely between the thorax and the abdomen. Genomic DNA was extracted from mosquito abdomens using ethanol precipitation method as described by Collins et al. [33]. One universal reverse primer and five animal-specific forward primers (human, cow, goat, pig, and dog) were used for amplification of cytochrome b gene, encoded in the mitochondrial genome to test for specific host blood meal origins using conventional PCR [35].

### Detection of Sporozoite Infectivity

The head and thorax of individual mosquitoes samples collected were used to detect the presence of *P. falciparum* sporozoites using enzyme-linked immunosorbent assays (ELISA) method as described by Wirtz et al. [36].

### Data analysis

Resting density of *Anopheline* mosquitoes was calculated as the number of female mosquitoes per trap/night for each trapping method. Analysis of variance (ANOVA) was used to compare malaria vector density between indoor and outdoor locations. Chi-square was used to test the difference in seasonal abundance and malaria vector species composition between resting locations (indoor and outdoor). Human blood index (HBI) was calculated as the proportion of blood-fed mosquito samples that had fed on humans to the total tested for blood meal origins [37]. Bovine, goat, dog, and pig blood indices were also calculated in a similar way. Mixed blood meals were included in the calculation of blood meal indices [38].

The sporozoite infection rate (IR) expressed as the proportion of mosquitoes positive for *Plasmodium* sporozoite was calculated by dividing the number of sporozoite positive mosquitoes by the total number of mosquitoes assayed. The frequency of the resistance allele was calculated using the Hardy-Weinberg equilibrium test for *kdr* genotypes. Data were analyzed using R software packages.

## Results

### Indoor and Outdoor *Anopheline* mosquito composition

A total of 2,706 and 860 female *Anopheline* mosquitoes were collected from Bungoma (highland site) and Kisian (lowland site) respectively during the study period. *Anopheles gambiae* s.l was the most abundant species accounting for 69.5% (1,880) in Bungoma and 90.8% (781) in Kisian followed by *An. funestus* 30.5% (826) and 9.2 % (79) respectively. Overall, the proportion of *Anopheles* species resting indoors was significantly higher by 82.4% than outdoor location 17.6% across the study sites (z = −8.47, p < 0.0001). The mean indoor resting density of *An. gambiae* s.l from both sites was significantly higher than outdoor resting density (*F*_1, 655_ =41.928, p < 0.0001). The mean indoor resting density for *An. funestus* was also higher than outdoor resting density (*F*_1, 655_ = 36.555, p < 0.0001) (Fig.2).

**Figure 1:**
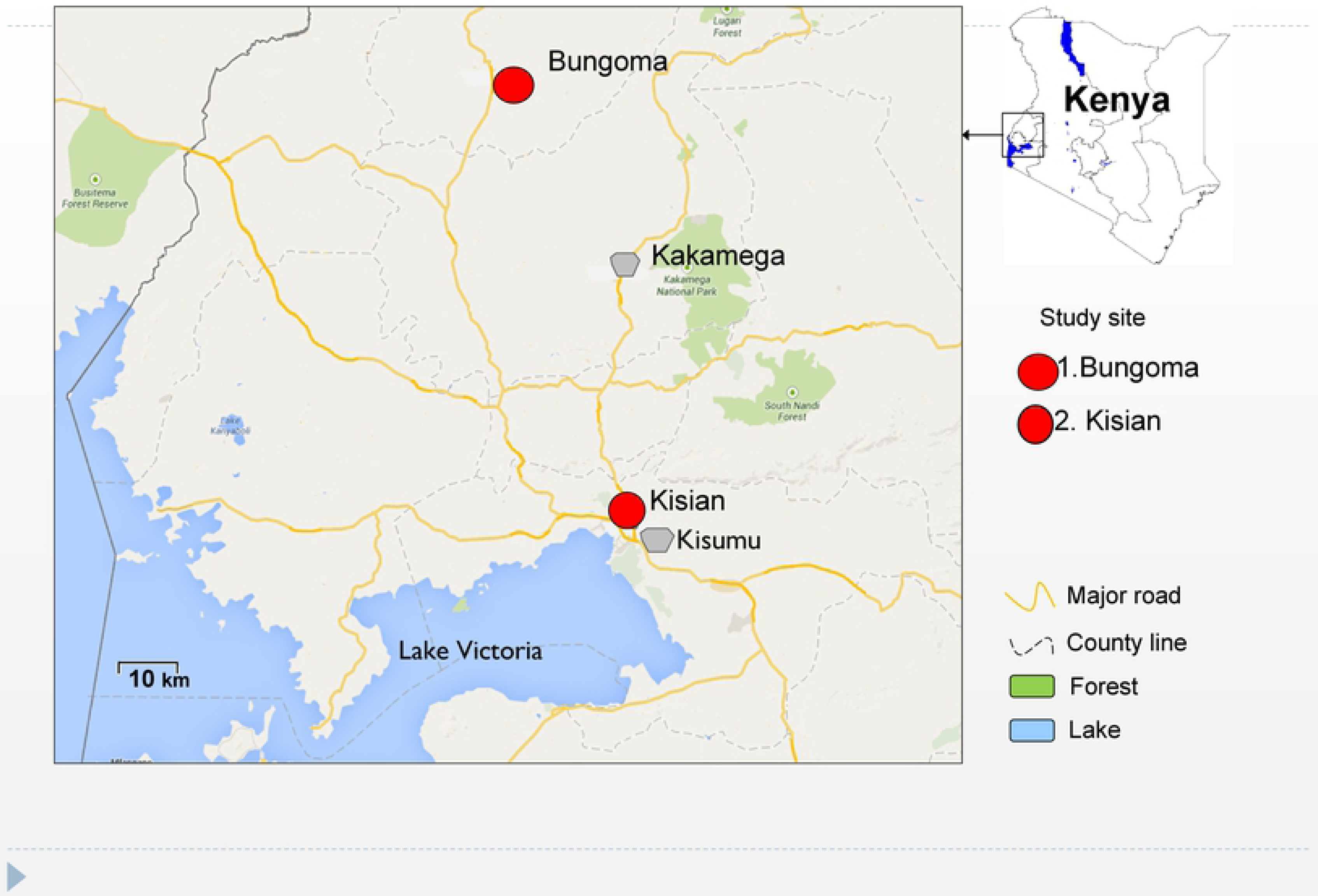
Map of the study sites in western Kenya

**Figure 2:**
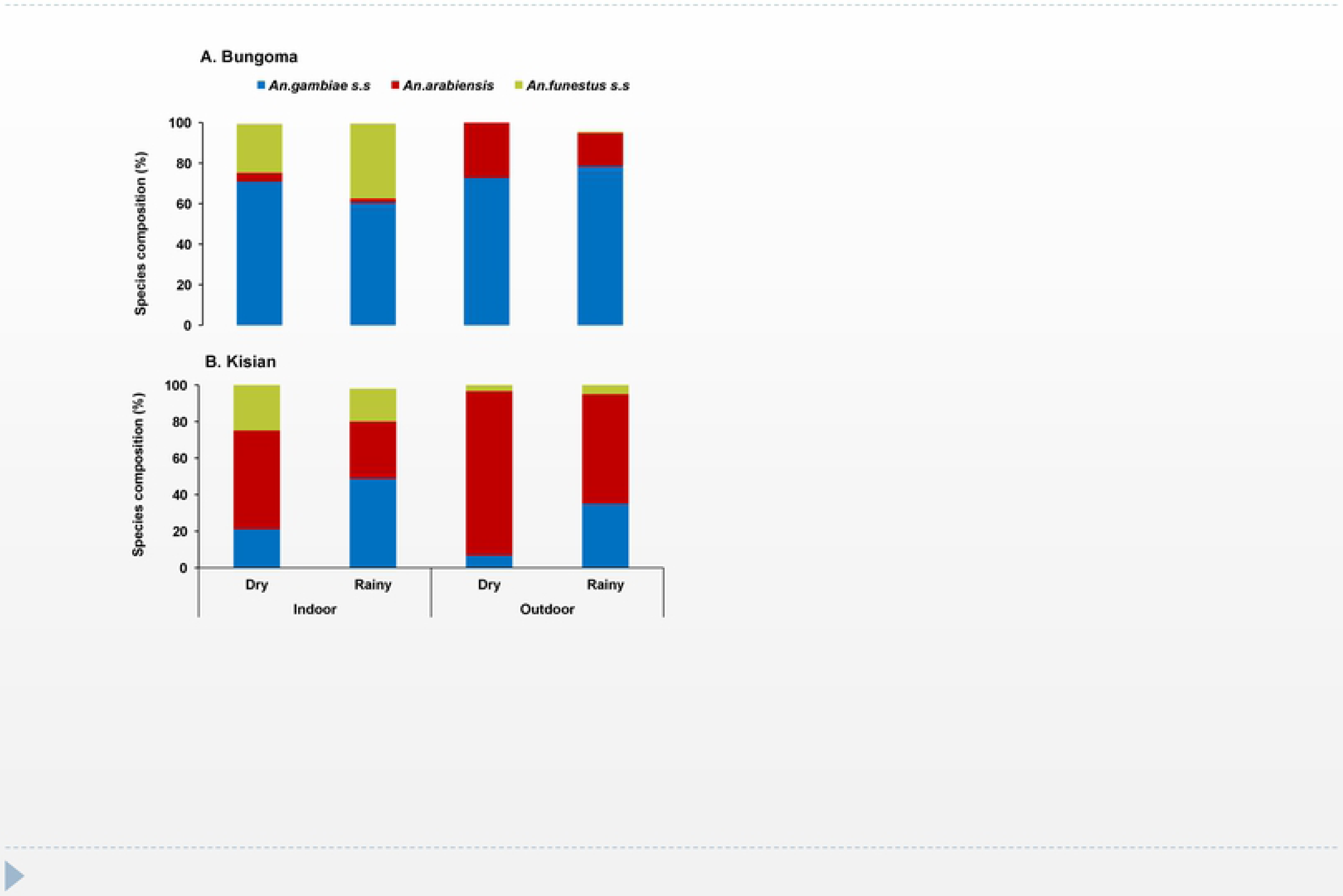
Indoor and outdoor resting density of female *Anopheles* mosquitoes collected per trapping method A: Highland site (Bungoma) and B: lowland site (Kisian), western Kenya. Error bars indicate 95% confidence intervals

For species identification, a sub-sample of 1,566 from both sites (1,172 *An. gambiae* s.l and 394 *An. funestus* s.l) were used to discriminate the sibling species. In Bungoma, *An. gambiae* and *An. arabiensis* accounted for 90.9% and 7.6% respectively. In Kisian, 60.2% were *An. arabiensis* and 38.9% as *An. gambiae.* All the *An. funestus* s.l assayed from the two study sites were all *An. funestus s.s* (hereafter *An. funestus*). Of the three vector species, *An. gambiae* (66.8%) was the predominant malaria vector in Bungoma followed by *An. funestus* (27.7%) and *An. arabiensis (*5.4%). In Kisian, *An. Arabiensis* (49.5%) was the most abundant vector species followed by *An. gambiae* (31.8%) and *An. funestus* (18.5%). There were more *An. gambiae* and *An. funestus* resting indoors than outdoors (80.7 vs 19.3% and 97.8 vs 2.2% respectively; X^2^ =122.96, *df* = 2, p < 0.0001). The proportion of *An. arabiensis* was higher outdoors than indoors (54.7 *vs* 45%). In Kisian, there were significantly more vectors resting indoors than outdoors (X^2^ =21.32, *df* = 2, p < 0.0001). There were more *An. gambiae* resting indoors (87.5%) than outdoors (12.5%). The densities of *An. arabiensis* (74.2 vs 25.8%) and *An. funestus* (96.0 vs 3.9%) also followed the same trend (Fig.3).

**Figure 3:**
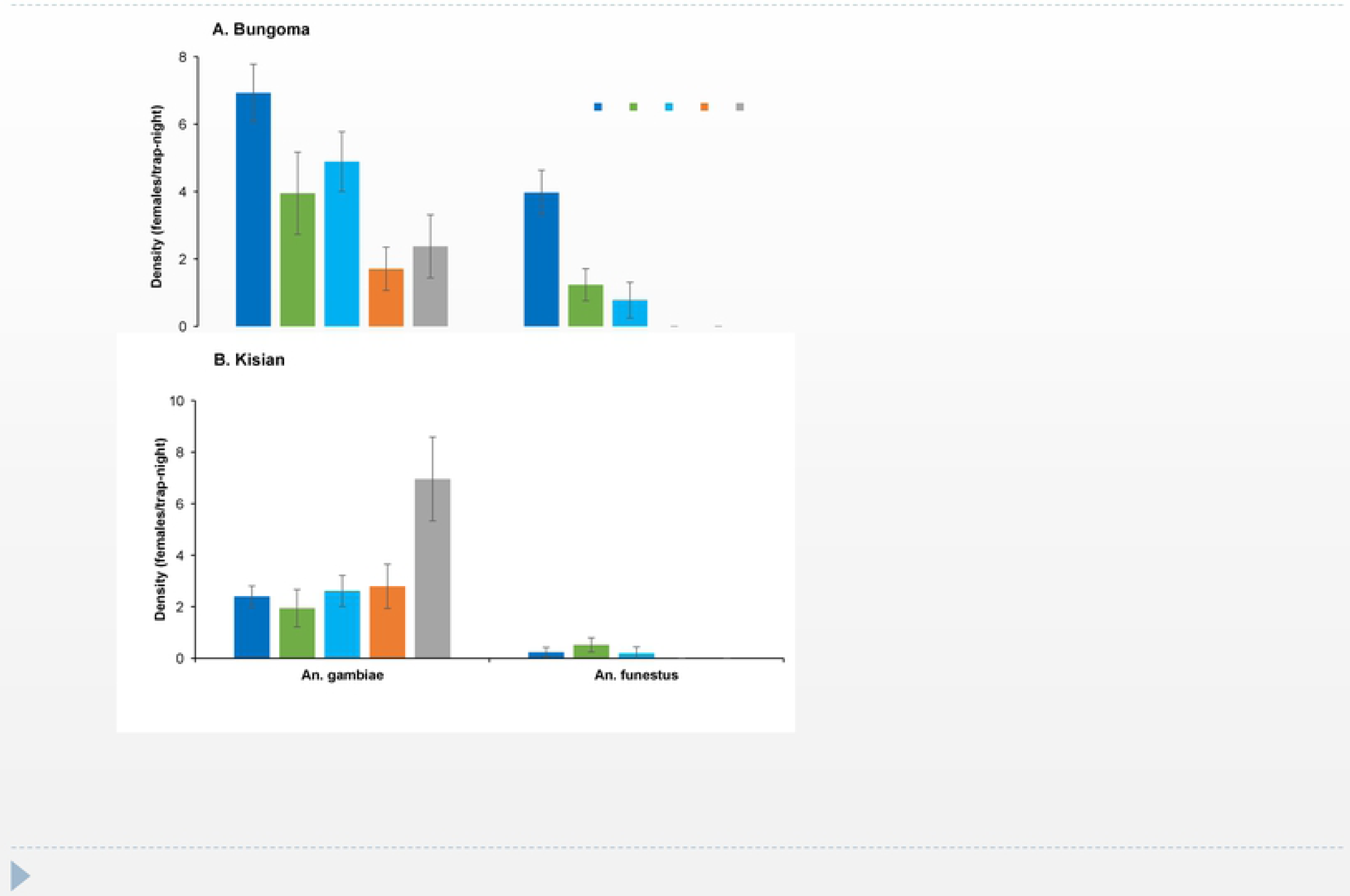
Seasonal abundance and *Anopheles* sibling species composition resting indoors and outdoors in A: Highland site (Bungoma) and B: lowland site (Kisian), western Kenya

In Bungoma, the seasonal abundance of *An. gambiae and An. funestus* species composition was higher during the rainy season 57% (352) vs 67% (211) than dry season 43 (266) vs 32.2% (100) indoors respectively (X^2^ =16.28, *df* = 2, p < 0.0003). However, *An. arabiensis* composition was higher during the dry season than rainy season indoors (63 vs 37%). There was no significant difference in *An. gambiae, An. arabiensis and An. funestus* species composition during the rainy season (78, 16.5 and 4.4% respectively) and dry season outdoors (*An. gambiae* 72.7% and *An. arabiensis*, 27.3%) in Bungoma (P>0.05). In contrast, in Kisian the overall seasonal prevalence of the three vector species composition was higher during the dry season indoors than rainy season (*An. arabiensis,* 68% (100) vs 32% (47), *An. funestus*, 63 (46) vs 37% (27); X^2^ =30.42, *df* = 2, p < 0.0001). However, the proportion of *An. gambiae s.s* was higher indoors during the rainy season 65.2% than dry season 34.8% (Fig.3).

### Phenotypic resistance

All the F1 mosquito populations tested from Bungoma and Kisian showed remarkable resistance to deltamethrin, with mortality rates ranging from 31.6% to 75.7% (Fig.4). High resistance levels were observed for F1 progeny of *Anopheles gambiae s.l* resting indoors (36.6%) than outdoors 65.5% in Bungoma. In Kisian the F1 progeny of *Anopheles gambiae s.l* resting indoors had lower mortality rates (66.6%) than outdoors (75.7%). Though the levels of deltamethrin resistance observed were higher for mosquitoes resting indoors compared to outdoors across the sites, there was no significant difference between the means (*F*_3_, _28_ =1.391, p < 0.266). Mortality rate for F1 progeny of *Anopheles funestus* resting indoors from Bungoma was 31.9%. Due to technical difficulties in raising *An. funestus* and the small numbers collected resting outdoors, susceptibility test was not done in Kisian and for the outdoor population from Bungoma. The *An. gambiae s.s* Kisumu susceptible laboratory strain showed 100% mortality.

**Figure 4:**
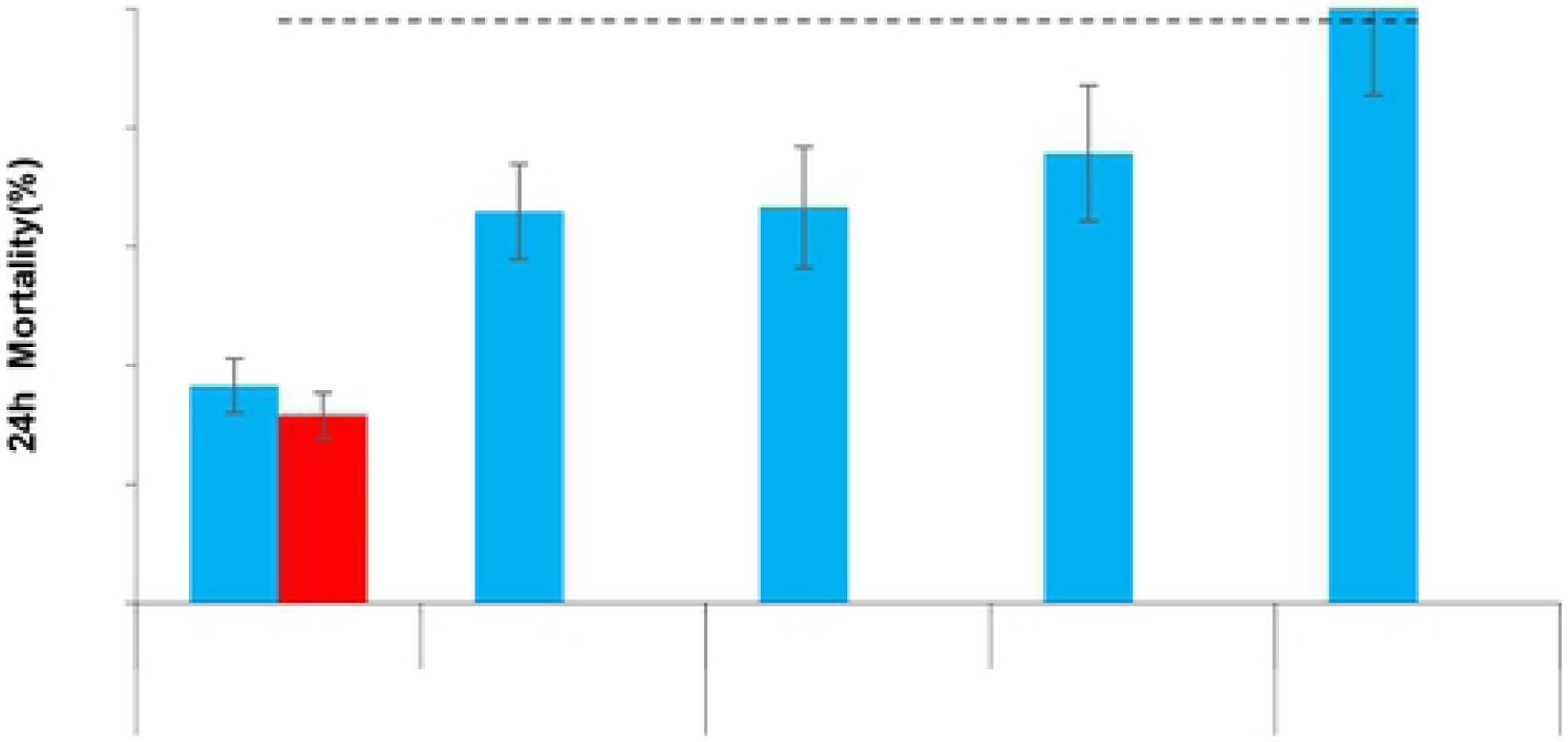
Mortality rates of indoor and outdoor resting *Anopheles gambiae* s.l and *An. funestus* F1 progeny in standard WHO tube bioassays after exposure to 0.05% deltamethrin test papers and 24 hr recovery period. Dotted lines represent upper (98%) and lower (90%) cut-offs for WHO classifications; values above the upper line indicate susceptibility and values below the lower red line indicate resistance (WHO, 2016). The vertical bar stands for standard error of the mean.

### Target site genotyping

In total 693 *Anopheles, gambiae* s.l samples were genotyped for the presence of Vgsc-1014S and 1014F mutations. In Bungoma, overall high frequency of Vgsc-1014S (87.7%) and 1014F (5.5%) was observed in An*. gambiae*, whereas only Vgsc-1014S was observed in *An. arabiensis* with a low frequency of 3.8%. The frequency of Vgsc-1014S and 1014F was high in indoor resting *An. gambiae* (89.7 and 9.2% respectively) than outdoor where only Vgsc-1014S was observed 84.6%. The Vgsc-1014S was the only kdr mutation observed in *An. arabiensis* resting indoors (10%) and was not detected in the outdoor resting collections (Table 1).

**Table 1:** Frequency of *Kdr* mutations in *An. gambiae s.s* and *An. arabiensis* populations of western Kenya

| Site | Species | Location | *An. gambiae* s.s |  |  | *An. arabiensis* |  |  |
| --- | --- | --- | --- | --- | --- | --- | --- | --- |
|  |  |  | N tested | 1014S | 1014F | N tested | 1014S | 1014F |
| Bungoma |  | Indoor | 195 | 89.7 | 9.2 | 20 | 10 | 0 |
|  |  | Outdoor | 133 | 84.6 | 0 | 32 | 0 | 0 |
|  | *An. gambiae* |  | 328 | 87.7 | 5.5 |  |  |  |
|  | *An. arabiensis* |  |  |  |  | 52 | 3.8 | 0 |
| Kisian |  | Indoor | 107 | 71.5 | 0.9 | 110 | 18.2 | 0 |
|  |  | Outdoor | 18 | 61.1 | 0 | 78 | 17.9 | 0 |
|  | *An. gambiae* |  | 125 | 70 | 0.8 |  |  |  |
|  | *An. arabiensis* |  |  |  |  | 188 | 18.1 | 0 |

In Kisian, the frequency of Vgsc-1014S in *An. gambiae* was 70% and that of 1014F was 0.8 %, whereas only Vgsc-1014S was observed in *An. arabiensis* with a frequency of 18.1 %. The frequency of Vgsc-1014S mutation was higher in *An. gambiae* resting indoors than outdoors (71.5 and 61.1% respectively). The same mutation was present in *An. arabiensis* collected from indoors 18.2% and outdoors 17.9%. The Vgsc-1014F was only observed in *An. gambiae* caught resting indoors at a low frequency of 0.8%.

### Blood meal sources

Blood-fed *An. gambiae* and *An. funestus* mosquito specimens were analysed for blood meal origins as shown in table 2. *An. funestus* was the most anthropophagic species and *An. arabiensis* the least from both sites. In Bungoma, the human blood index (HBI) for *An. gambiae*, *An. arabiensis* and *An. funestus* were 64.7, 25 and 74.8% respectively. The HBI for *An. gambiae* resting indoors was higher (54.3%) than outdoors (33.3%), and the HBI of *An. arabiensis* resting indoor (25.0%) was higher than for outdoor (12.5%). *An. funestus* was highly anthropophilic with HBI of 64.8% from indoors and 75% outdoors.

**Table 2:** Blood meal origins of *Anopheline* mosquitoes collected from indoor and outdoor in Bungoma and Kisian, western Kenya

| Site | Blood-meal origins | *An.gambiae s.s* |  | *An. arabiensis* |  | *An. funestus* |  |
| --- | --- | --- | --- | --- | --- | --- | --- |
|  |  | Indoor | Outdoor | Indoor | Outdoor | Indoor | Outdoor |
| Bungoma | **Number tested** | **422** | **39** | **24** | **8** | **111** | **4** |
|  | Human | 229 (54.3) | 13 (33.3) | 6 (25.0) | 1 (12.5) | 72 (64.9) | 3 (75.0) |
|  | Bovine | 60 (14.2) | 9 (23.1) | 16 (66.7) | 3 (37.5) | 14 (12.6) | 1 (25.0) |
|  | Human+Bovine | 31 (7.3) | 4 (10.3) | 0 | 1 (12.5) | 9 (8.1) | 0 |
|  | Other | 24 (5.7) | 4 (1.0) | 2 (8.3) | 1 (12.5) | 3 (2.7) | 0 |
|  | Un-identified | 78 (18.5) | 9 (23.1) | 0 | 2 (25.0) | 13 (11.7) | 0 |
|  | **HBI** | **64.7** | **46.2** | **25.0** | **37.5** | **74.8** | **75.0** |
|  | BBI | 22.2 | 35.8 | 66.7 | 62.5 | 20.7 | 25.0 |
| Kisian | **Number tested** | **77** | **3** | **114** | **32** | **23** | **0** |
|  | Human | 42 (54.5) | 2 (66.7) | 6 (5.3) | 0 | 18 (78.4) | 0 |
|  | Bovine | 26 (33.8) | 0 | 104 (91.2) | 29 (90.6) | 4 (17.4) | 0 |
|  | Human+Bovine | 3 (3.9) | 1 (33.3) | 2 (1.8) | 1 (3.13) | 1 (4.3) | 0 |
|  | Other | 4 (5.2) | 0 | 1 (0.9) | 2 (6.3) | 1 (4.3) |  |
|  | Un-identified | 2 (2.6) | 0 | 1 (0.9) | 0 | 0 | 0 |
|  | **HBI** | **59.7** | **100** | **7.0** | **6.3** | **82.7** | **0** |
|  | BBI | 40.2 | 33.3 | 93.0 | 96.9 | 21.7 | 0 |

Others: dog, goat, pig, or multi-source excluding human+bovine
HBI, human blood Index; BBI, bovine blood index

In Kisian, the overall HBI of *An. gambiae*, *An. arabiensis* and *An. funestus* was 59.7, 7 and 82.7% respectively. The HBI for *An. gambiae* resting outdoors was higher (100%) than indoors (59.7%). The HBI for indoor and outdoor resting *An. arabiensis* was 7 vs 6.25%. The HBI of *An. funestus* resting indoors was 82.4%. None of the *An. funestus* from outdoor collections was positive for a blood meal test.

In Bungoma, the overall Bovine blood index (BBI) for *An. gambiae*, *An. arabiensis* and *An. funestus* were 22.2, 66.7 and 20.7% respectively. The BBI for *An. gambiae* resting indoors were lower (14.2%) than outdoors (23.1%). *An. arabiensis* caught resting indoors had higher BBI (66.7%) than outdoors (37.5%). The BBI of *An. funestus* resting indoors and outdoors was 12.6 and 25% respectively. In Kisian, the overall BBI for *An. gambiae*, *An. arabiensis,* and *An. funestus* was 40.2, 93.0 and 21.7% respectively. The BBI of *An. gambiae* resting indoor was 33.8% and none tested positive for outdoors. *An. arabiensis* collected from both indoors and outdoors were highly zoophilic with BBI of 93% and 96.9% respectively. The BBI of *An. funestus* resting indoors was 17.4%. The blood meal indices for other hosts (goat, pig, and dog) were <10% from both sites (Table 2).

### Sporozoites infection rates

A total of 1,517 samples comprising of 1,156 *An. gambiae* s.l and 361 *An. funestus* specimens were tested for *Plasmodium falciparum* Circumsporozoite (CSP) (Table 3). Ninety-one samples (90 Bungoma and 1 Kisian) tested positive giving an overall infection rate of 7.8% in Bungoma and 0.3% in Kisian. Overall, the sporozoite rate was higher indoors (8.6%) than outdoors (4.2%) in Bungoma, whereas in Kisian the sporozoite rate was 0.3% indoors. None of the samples collected outdoors in Kisian tested positive. The overall sporozoite rate for *An. funestus* collected from Bungoma was 10.6%. *An. funestus* resting indoors had a sporozoite rate of 11%. None of *An. funestus* collected outdoors was positive. The sporozoite rate for indoor resting *An. gambiae* was 7.6% and outdoors at 4.7 %. For *An. arabiensis,* the overall sporozoite rate was 3.13% with those resting indoors and outdoors having sporozoite rates of 3.4 and 2.9 % respectively. The overall entomological inoculation rates (EIRs) of the three vector species from indoor resting collections and outdoor was 66 and 10.4 infective bites/person/year respectively. In Kisian, only 1 (0.9%) *An. gambiae* collected from indoor was CSP positive. No CSP positives were detected in *An. arabiensis* and *An. funestus* resting indoors and outdoors in Kisian. The overall EIR from indoor collections was 1.38 infective bites/person/year.

**Table 3:** Sporozoite rates of *Anopheles* mosquitoes from indoor and outdoor collections in Bungoma and Kisian, western Kenya

| Site | Season | Location | *An. gambiae s.s* |  | *An. arabiensis* |  | *An. funestus* |  |
| --- | --- | --- | --- | --- | --- | --- | --- | --- |
|  |  |  | N tested | Pf +ve | N tested | Pf +ve | N tested | Pf +ve |
| Bungoma |  | Indoor | 618 | 47(7.6) | 27 | 1(3.7) | 311 | 34(10.9) |
|  |  | Outdoor | 148 | 7(4.7) | 35 | 1(2.6) | 7 | 0 |
|  | Dry |  | 290 | 21(7.2) | 26 | 0 | 100 | 18(18.0) |
|  | Rainy |  | 476 | 33(6.9) | 36 | 2(5.6) | 218 | 17(7.9) |
| Kisian |  | Indoor | 112 | 1(0.9) | 147 | 0 | 73 | 0 |
|  |  | Outdoor | 16 | 0 | 54 | 0 | 3 | 0 |
|  | Dry |  | 41 | 1(2.4) | 127 | 0 | 47 | 0 |
|  | Rainy |  | 87 | 0 | 74 | 0 | 29 | 0 |

Pf, *Plasmodium falciparum*
+ve, Positive

In Bungoma, the overall seasonal infective rate was high during the dry season (18%) than in the rainy season (7.6%) for *An. funestus*. For *Anopheles gambiae*, the seasonal infective rate was 7.2% during the dry season and 7.06% during the rainy season. *Anopheles arabiensis* seasonal infective rate was only detected during the rainy season (5.6%) (Table 3). In Kisian, the seasonal infective rate of *An. gambiae* was 2.4 % during the dry season.

## Discussion

Given the current widespread of pyrethroid insecticide resistance in western Kenya [16,18,39], little is known about the behavioural response of these malaria vector population to the wide use of LLINs. Evidence has shown that successful malaria elimination strategies require vector control interventions that target the changing vector behaviour [40]. Overall, this study investigated the behavioral heterogeneity of malaria vectors for resting behavior, feeding choices and infection rates in the context of increased use of LLINs. The study revealed high indoor densities and high frequency of kdr mutations indoors compared to outdoors, leading to increased vector-human contact and ongoing malaria transmission in the region despite the high coverage and usage of LLINs.

Of the three major malaria vectors in western Kenya, higher proportions of *An. gambiae* and *An. funestus* were abundantly caught resting inside houses in both Bungoma (highland site) and Kisian (lowland site), with the dominance of *An. gambiae* at Bungoma, whereas a considerable number of *An. arabiensis* was found resting outdoors in both sites and was abundant at Kisian. This distribution and variation in the relative frequency and behavioral patterns of the three malaria vectors according to ecological setting is consistent with earlier reports [15,19,41–43]. The high indoor resting behavior of *An. gambiae* and *An. funestus* might be attributed to the reduced susceptibility of these vectors to insecticides used for vector control. This may increase human-vector contact and lead to high malaria transmission indoors. *An. arabiensis* has been known to be more flexible in terms of feeding and resting behavior at different locations, with some of these variations explained by the historical use of insecticides [22, 44]. Its potential to rest outside human dwellings increases its chance to survive the current indoor intervention, hence establishing extra-domiciliary malaria transmission. It is worth noting that, the proportion of *An. arabiensis* was higher during the dry season in Kisian (lowland)*. An. arabiensis* has been shown to survive best in the lowland sites where temperatures are higher and relative humidity is lower than the highlands[45], this could favor the survival of this species in the region.

This study observed a high frequency of Vgsc-1014S and Vgsc-1014F from indoors resting *An. gambiae* than outdoors, from all sites. These findings are in agreement with previous studies from the same region that observed increased Kdr frequency in *An. gambiae* s.l [15,46,47]. The Vgsc-1014F mutation was only detected in *An. gambiae* resting indoors, this mutation was absent in the mosquitoes collected outdoors. Additionally, lower mortality rates were observed in F1 progeny of the indoor resting mosquitoes exposed to deltamethrin than outdoors. This finding suggests that malaria vectors with high resistance levels might tend to increase their feeding on humans by remaining indoors, regardless of the presence of indoor vector control interventions. Initial studies have observed the occurrence of Vgsc-1014F at low frequencies in East Africa including Kenya [39,48,49]. The presence of kdr mutations at Bungoma where it was first detected in Kenya in *An. gambiae* [15] and now at Kisian in the lowlands indicate the widespread occurrence of the mutations among the *An. gambiae* population. The presence of Vgsc-1014F in indoor resting mosquitoes may be of the particular concern given that the mutation has been found to be strongly associated with pyrethroid resistance than *Vgsc-1014S* which has been observed to be a weaker kdr mutation [50]. The increased resistance level could be a result of selection force due to the widespread use of LLINs leading to the selection of resistant strains [5]. Some studies have shown a relationship between the rapid spread of kdr alleles with the widespread use of LLINs [51–53]. The extensive use of agriculture insecticides may also contribute to the occurrence of new mutations to existing insecticides [41,54–57].

Even though we observed high frequencies of Vgsc-1014S in *An. gambiae,* the allele was at low frequency in *An. arabiensis,* with a higher frequency of Vgsc-1014S, detected for indoor resting individuals than outdoors. Most recently, the presence of kdr mutations in *An. arabiensis* from the lowlands of western Kenya has been reported [16, 58]. The low kdr frequency observed in *An. arabiensis* could be due to the reduced insecticide selection pressure imposed on them as they resort to feed and rest outdoors in the absence of insecticides, unlike *An. gambiae* that feeds and rests indoor. The frequent kdr mutations, behavioural resilience and an increased proportion of *An. arabiensis* resting outdoors could all raise further concerns on the future utility of the current indoor interventions.

*Anopheles gambiae* and *An. funestus* have been known to be efficient in malaria transmission due to their anthropophagic and endophilic behaviour [19,59,60]. Blood meal analysis results showed a large proportion of *An. gambiae* and *An. funestus* that are resting indoors, preferred feeding on humans than animals. The high HBI observed in the two species resting indoors could be partly explained by increased insecticide resistance observed in this population, as the current indoor-based interventions could have a minimal effect on their daily activities indoors. This human-host choice and higher indoor resting proportions of *An. funestus* poses a great concern in malaria elimination efforts due to its efficiencies in transmitting malaria. *An. arabiensis* resting indoors and outdoors both took a larger proportion of blood meals from cattle than did either *An. gambiae* or *An. funestus.*The observed zoophilic behavior could be attributed to the increased susceptibility of this vector to the current indoor intervention.

Sporozoite infection rates were high in *An. funestus* and *An. gambiae* collected from Bungoma, with *An. funestus* showing considerably higher sporozoite rates than the other species. These findings confirm earlier reports from other regions in western Kenya that have documented the re-emergence of *An. funestus* and its role in malaria transmission [19, 60]. The majority of malaria infection rates were higher for vectors collected indoors than outdoors, suggesting ongoing malaria transmission regardless of the use of LLINs. The probable explanation for high sporozoite rates indoors could be due to inconsistency usage of LLINs [61] and increased insecticide resistance of malaria vectors to available interventions as the vectors spend more time indoors than outdoors. This implies that the insecticide-treated materials used indoors to reduce human-vector contact might be less effective in repelling and killing malaria vectors. Some studies have linked the rebound of malaria with insecticide resistance after high coverage of LLINs [51,53,62]. The level of phenotypic resistance in *An. gambiae s.l* and *An. funestus* together with the frequency of insecticide resistance markers observed in this study confirms the above assertion. The observed sporozoite infection rates outdoors might be attributed to changing in the biting behaviour of malaria vectors as some vectors could be feeding on humans when they are active and unprotected outdoors [63]. This need for complementary tools such as repellents [64]for personal protection or larval control[65] to suppress malaria transmission outdoors as the current indoor tools alone may seem to be insufficient to drive malaria infections. *An. gambiae* and *An. arabiensis* were responsible for outdoor malaria transmission, with *An. arabiensis* showing low positivity rates than *An. gambiae.* This could be partly due to the zoophilic nature of *An. arabiensis* [22]. Despite the low sporozoite rates of *An. arabiensis* reported in this study, its importance in outdoor malaria transmission should not be assumed due to its opportunistic behavior as the vector could continue with the residual transmission outdoors in the region, which is a major threat to effective malaria vector control. The study recorded low sporozoite rates in Kisian (lowland). This could be due to the dominance of *An. arabiensis* in the region.

## Conclusion

The study shows high densities of insecticide-resistant *An. gambiae* and *An. funestus* resting indoors despite the use of indoor interventions with the increasing importance of *An. funestus* in sustaining malaria transmission in western Kenya highlands. *An. arabiensis* were more outdoors than indoors. This behavioural plasticity increases its survival and potential in continuing residual transmission after the main endophilic and endophagic vectors have been reduced by the interventions. The Vgsc-1014S and Vgsc-1014F mutations were observed at high frequencies in *An. gambiae* resting indoors. This calls for further screening of other resistance mutations in this population for better resistance management. Sporozoite rates were higher indoors than outdoors, showing that transmission occurs more indoors than outdoors in these sites. Insecticide resistance management strategies and/or new vector control interventions that may not insecticide based are needed in western Kenya to reduce malaria transmission.

## Abbreviations

F1: first generation offspring
PCR: polymerase chain reaction
ELISA: enzyme-linked immunosorbent assay
Vgsc: voltage-gated sodium channel
HBI: human blood index
LLINs: long-lasting insecticidal nets
IRS: indoor residual spray
PSC: pyrethrum spray catch
KDR: Knockdown resistant gene
BBI: bovine blood index
CSP: circumsporozoite protein
EIR: entomological inoculation rate

## Ethical Considerations

The institutional review board of Kenya Medical Research Institute granted ethical review and approval (Ref: KEMRI/SERU/CGHR/085/3434). Prior to the commencement of data collection, a detailed explanation of the aims, study procedures, risks and benefits were provided to community leaders and participants of each study site. Informed consent was obtained from the household heads. Participation was voluntary and household heads were free to withdraw from the study in case of any inconvenience. The permission to publish this study was granted by the director of Kenya Medical Research Institute.

## Acknowledgments

The authors wish to thank the villagers and community leaders in Bungoma and Kisian for their permission to collect mosquitoes in their houses. In particular, we thank the community leaders. We acknowledge the Entomology Laboratory at Kenya Medical Research Institute, Kisumu for providing technical and laboratory space for the study.

## Authors’ contribution

MM, EO, GY, and YAA conceived and designed the experiments MM participated in data collection, performed laboratory work, MM, JK and GZ data analysis and drafted the manuscript. EO, FA, SM, AK, GY, and YAA, supervised data collection. All authors have read and approved the final manuscript.

## Availability of data and materials

The datasets used for the current study are available from the corresponding author on reasonable request.

